# Architectural RNA is required for heterochromatin organization

**DOI:** 10.1101/784835

**Authors:** Jitendra Thakur, He Fang, Trizia Llagas, Christine M. Disteche, Steven Henikoff

## Abstract

In addition to its known roles in protein synthesis and enzyme catalysis, RNA has been proposed to stabilize higher-order chromatin structure. To distinguish presumed architectural roles of RNA from other functions, we applied a ribonuclease digestion strategy to our CUT&RUN in situ chromatin profiling method (CUT&RUN.RNase). We find that depletion of RNA compromises association of the murine nucleolar protein Nucleophosmin with pericentric heterochromatin and alters the chromatin environment of CCCTC-binding factor (CTCF) bound regions. Strikingly, we find that RNA maintains the integrity of both constitutive (H3K9me3 marked) and facultative (H3K27me3 marked) heterochromatic regions as compact domains, but only moderately stabilizes euchromatin. To establish the specificity of heterochromatin stabilization by RNA, we performed CUT&RUN on cells deleted for the *Firre* long non-coding RNA and observed disruption of H3K27me3 domains on several chromosomes. We conclude that RNA maintains local and global chromatin organization by acting as a structural scaffold for heterochromatic domains.

## Introduction

Eukaryotic genomic DNA is packaged with histone proteins into nucleosomes, which serve as the repeating structural unit of chromatin (Zhou et al., 2019). Two forms of chromatin, heterochromatin and euchromatin, were originally defined respectively as condensed (strongly staining) and decondensed (weakly staining) chromatin states (Heitz 1928). Euchromatin assembles on relatively gene-rich DNA and localizes preferentially to the interior of the nucleus (Bártová et al., 2008; Burke and Stewart, 2014). Euchromatin is enriched for RNA Polymerase II and is marked by active histone modifications such as trimethylation of lysine 4 in histone H3 (H3K4me3), whereas heterochromatin assembles on gene-poor DNA and is marked by repressive histone modifications. Constitutive heterochromatin is marked by trimethylation of H3 lysine 9 (H3K9me3) and assembles mostly on repetitive DNA, whereas facultative heterochromatin is marked by trimethylation of H3 lysine 27 (H3K27me3) and assembles mostly on developmentally regulated loci (Cheutin and Cavalli, 2012; Pinheiro and Heard, 2017; Politz et al., 2013). H3K27me3 and H3K9me3 modifications are recognized by the chromodomains of the Polycomb repressive complex and heterochromatin protein 1 (HP1) respectively, which then recruit other chromatin modifiers to propagate the heterochromatic state (Bannister et al., 2001; Lachner et al., 2001).

In addition to DNA-binding chromatin modifiers, RNA has emerged as an important regulator of heterochromatin formation. In fission yeast, small RNA transcribed from pericentric DNA repeats are critical for RNAi-mediated H3K9me3 heterochromatin formation (Grewal, 2010). In mammals, long non-coding RNAs (lncRNAs) are essential for the onset of silent heterochromatin during X-chromosome inactivation (*Xist* and *Tsix* lncRNAs) and genome imprinting (*Kcnq1ot1* and *Air* lncRNAs) (Nagano et al., 2008; Pandey et al., 2008; Pinheiro and Heard, 2017; Verdel et al., 2004; Zhao et al., 2008). Chromatin modifiers physically interact with several RNAs (Engreitz et al., 2016; Pandey et al., 2008; Prensner et al., 2013; Rinn et al., 2007; Yang et al., 2015; Yap et al., 2010), suggesting a widespread role of RNA in chromatin organization. In addition to the indirect regulatory role in which RNA contributes to chromatin organization by recruiting chromatin modifiers, a direct architectural role of RNA in chromatin organization has been proposed (Nickerson et al., 1989). Cytological visualization of Ribonuclease A (RNase A) treated cells has shown loss of a specific H3K9me3 branched epitope (which comprises a small fraction of pericentric heterochromatin), leaving the rest of the heterochromatin intact (Maison et al., 2002). Genome-wide chromatin conformation capture (Hi-C) assays in RNase A treated cells suggests that although the overall genome organization and the higher-order chromatin organization such as topologically associated domains (TADs) remain unaffected, a subtle change in long-range interactions within heterochromatic, gene-poor, and silent genomic compartments is observed upon RNA depletion (Barutcu et al., 2019). However, the low resolution of both cytology and Hi-C methods limits their use in deciphering the involvement of RNA in chromatin structure.

Here, we ask whether the physical presence of RNA affects chromatin organization by combining our high-resolution in situ genomic profiling method, Cleavage Under Targets and Release Using Nuclease (CUT&RUN) (Skene and Henikoff, 2017), with RNase A treatment. We show that the presence of architectural RNA is necessary for the maintenance of the local chromatin environment and for the structural integrity of heterochromatic regions. We demonstrate the specificity of RNA-heterochromatin interactions by showing that the Firre lncRNA stabilizes facultative heterochromatin on multiple chromosomes.

## Results

CUT&RUN.RNase probes the requirement of architectural RNA in perinucleolar chromatin structure

Chromatin profiling methods such as Chromatin Immunoprecipitation (ChIP) (Park, 2009), require chromatin to be fragmented before the antibody binding step and thus lose native 3-D conformation across the targeted epitope. In contrast, CUT&RUN is performed using intact permeabilized cells, thereby preserving the structure of chromatin and other macromolecular ensembles around the chromatin feature of interest. In CUT&RUN, a fusion of Micrococcal Nuclease and protein-A (pA-MN) is targeted to chromatin via an antibody. Subsequently, pA-MN is activated in the presence of Ca++ ions to catalyze digestion at targeted loci. We modified CUT&RUN to CUT&RUN. RNase by digesting total RNA of permeabilized cells by adding RNase A before the antibody binding step, to rapidly digest total RNA of permeabilized cells and observe its effect upon chromatin structure (Schematic in Figure 1A). If the physical presence of RNA is essential for the native chromatin conformation, then removal of RNA from cells should alter CUT&RUN signals. Because RNA is removed in situ from intact cells, CUT&RUN.RNase detects direct structural interactions of RNA with chromatin, whether or not the RNAs also have indirect structural roles.

**Figure 1.**
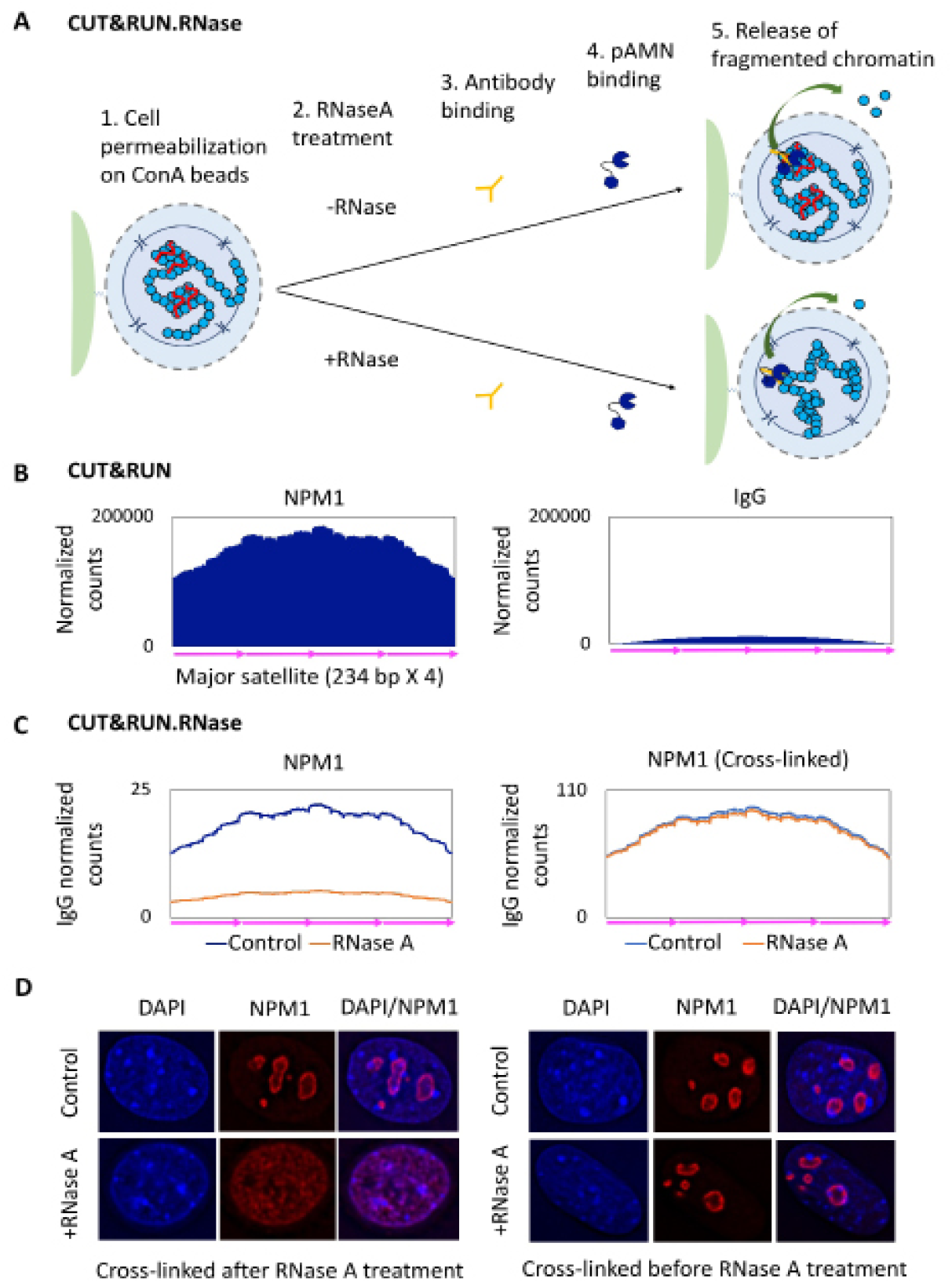
CUT&RUN.RNase identifies RNA-chromatin interactions. A) Schematic of the CUT&RUN.RNase method B) Average profiles of normalized NPM1 and IgG CUT&RUN on a concatenated major satellite consensus sequence (234 bp repeat unit) revealed a clear enrichment of Nucleophosmin (NPM1) on pericentric heterochromatin relative to IgG. C) The loss of NPM1 signals on pericentric major satellites in NPM1 CUT&RNA.RNase (Left). Cross-linking retains NPM1 on the pericentric heterochromatin in CUT&RNA.RNase (Right). D) Cytological visualization of NPM1 under indicated conditions.

To determine whether CUT&RUN.RNase can be used to detect the dependence of chromatin integrity on its direct contacts with RNA, we applied CUT&RUN. RNase to the chromatin surrounding the nucleolus, a major hub of nuclear RNA, in Patski murine embryonic kidney fibroblast cells. The outer part of the nucleolus, called the granular component (GC), is in direct contact with nucleoplasm and is occupied by several proteins including Nucleophosmin (NPM1) (Boisvert et al., 2007; Lindström, 2011). Integration of NPM1 into the nucleolus is dependent on its multivalent interactions with RNA and proteins containing arginine-rich linear motifs (Mitrea et al., 2016). Furthermore, loss of NPM1 triggers rearrangement of perinucleolar heterochromatin (Holmberg Olausson et al., 2014; Murano et al., 2008), suggesting direct interactions between NPM1 and heterochromatin. Perinucleolar heterochromatin in mouse cells is mostly pericentric and consists of major satellite DNA (Almouzni and Probst, 2011). We profiled NPM1 using CUT&RUN and detected a clear enrichment of NPM1 on major satellite DNA in Patski cells (Figure 1B, Supplementary Figure 1). Next, to determine if NPM1 interactions with heterochromatin are dependent on the physical presence of RNA, we performed NPM1 CUT&RUN.RNase profiling. To compare RNase-treated to untreated cells, we calibrated datasets using an IgG negative control performed on the same RNase-treated cells. We observed dramatic loss of NPM1 signals on major satellites upon RNase A treatment (Figure 1C), which reveals that NPM1 interactions with pericentric chromatin are dependent on the physical presence of RNA.

If the decrease in NPM1 signals on pericentric heterochromatin is due to RNA-dependent anchoring, and is not an artifact of RNase A treatment, cross-linking prior to in situ removal of RNA should prevent the loss of NPM1 from pericentric heterochromatin at the nucleolar periphery. Indeed, when we cross-linked the cells prior to CUT&RUN. RNase, we observed complete rescue of NPM1 signals in RNase A treated cells, strongly suggesting that the reduction in CUT&RUN signals results from structural disintegration (Figure 1B). Cytological visualization of NPM1 in control cells revealed a bright NPM1 ring around each nucleolus. When RNase A treatment was performed before formaldehyde cross-linking, the NPM1 ring completely collapsed, whereas when RNase A treatment was performed after cross-linking the cells, NPM1 rings stayed intact and resembled those in untreated cells (Figure 1D). Together these results suggest that the RNA is necessary for NPM1-pericentric heterochromatin interactions at the nucleolar periphery and that CUT&RUN can be used to probe the requirement of the physical presence of RNA in chromatin integrity/organization.

### RNA depletion disrupts the local chromatin environment around CTCF binding sites

The major genome architectural protein CCCTC-binding factor (CTCF) binds either directly to its target CTCF motif or indirectly via long range interactions with sites that are in physical proximity (Phillips and Corces, 2009; Skene and Henikoff, 2017). CTCF also shapes local chromatin by positioning multiple nucleosomes flanking its binding site (Fu et al., 2008). These CTCF interactions are dependent on its RNA binding domain, suggesting a critical role for RNA in CTCF-mediated chromatin organization (Anders et al., 2019, Saldana-Meyer et al., 2019). It remains unclear whether the contribution of RNA in maintaining CTCF-chromatin interactions is due to direct RNA binding or to an indirect regulatolatory role. To distinguish these possibilities, we profiled CTCF using CUT&RUN.RNase. This revealed a 2.8-fold decrease in CTCF signal (Figure 2A-B). Mapping of CTCF CUT&RUN sequencing reads results in two fragment size classes - smaller fragments peaking at ~100 bp corresponding to direct CTCF footprints and larger fragments peaking at ~150 bp resulting from the action of pA-MN within linker regions of flanking nucleosomes (Skene and Henikoff, 2017). Fragment length analysis revealed a reduction in CTCF signals upon RNA depletion in both small and large CTCF fragments (Figure 2A-B). We observed that the fraction of large CUT&RUN fragments mapped to CTCF sites decreased significantly upon RNA depletion, reflecting a greater decrease in cleavages within the linker regions of nucleosomes flanking CTCF as compared to those adjacent to direct CTCF binding sites in mouse embryonic kidney cells (Figure 2C). Similarly, CTCF CUT&RUN.RNase in HeLa cells resulted in decreased CTCF signals on both direct (containing a CTCF-specific binding DNA motif) and indirect 3-D contact sites (lacking a CTCF motif) (Supplementary Figure 2). These results reveal that RNA depletion reduces the digestion and release of flanking chromatin during CUT&RUN, suggesting that the presence of RNA helps to maintain chromatin compaction around CTCF sites.

**Figure 2.**
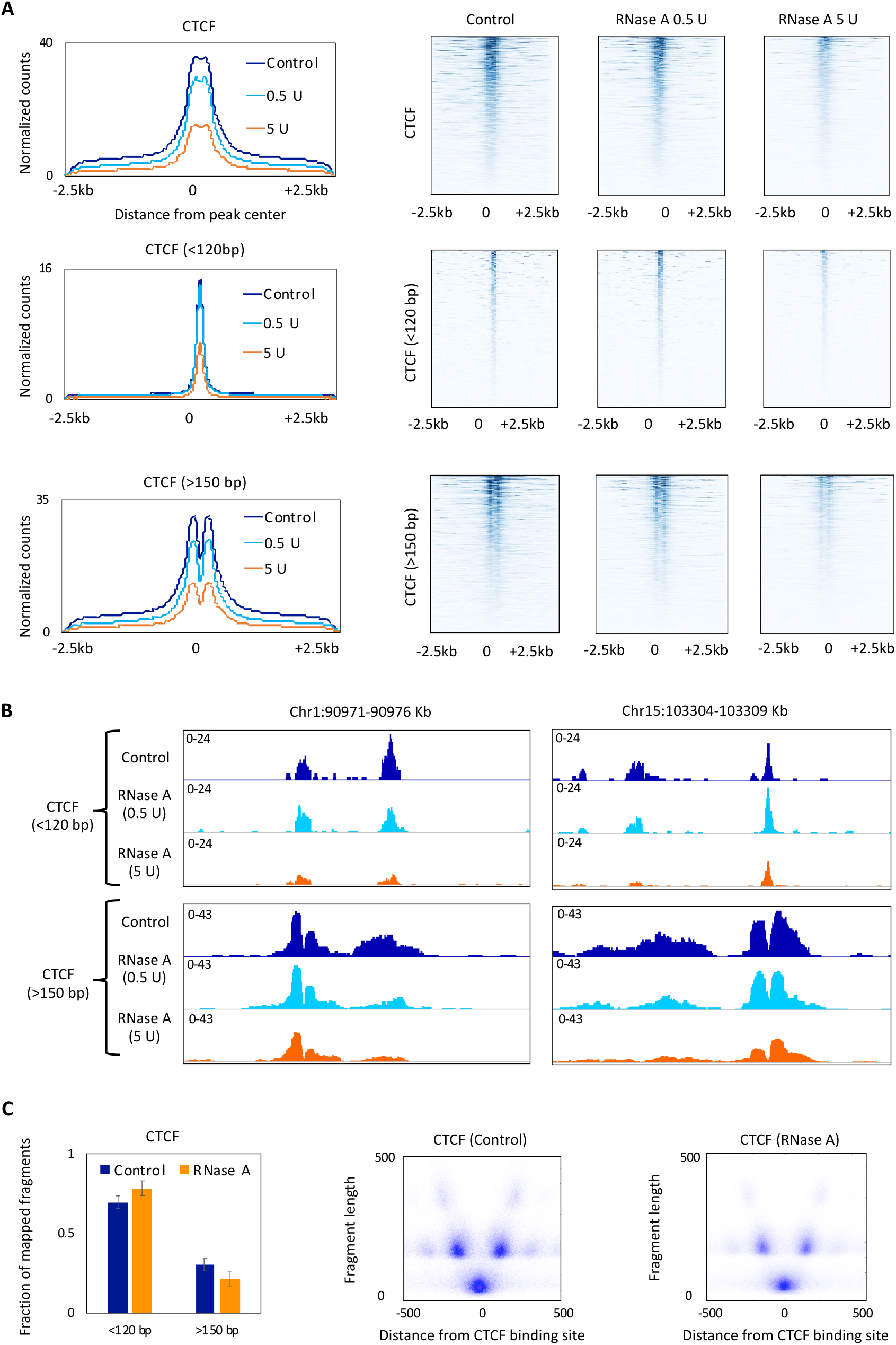
CUT&RUN.RNase reveals an altered local chromatin environment at CTCF sites upon RNA depletion. A) CTCF CUT&RUN.RNAse signals for all, small (< 120 bp) and large (> 150 bp) fragments in control cells and cells treated with RNase A treated cells (0.5 and 5 U RNase A per million cells). Genome-wide average profiles (Left) and heatmaps (Right) were generated on a 5 kb region across annotated CTCF sites. B) CTCF CUT&RUN.RNase tracks across two representative 5-kb genomic regions. C) Fraction of small and large CTCF fragments in control and RNase A treated cells (Left). V-plot of CTCF on 600-bp region spanning CTCF sites (Right).

### RNA depletion disrupts the structural integrity of heterochromatin

Next, to determine if the presence of RNA maintains the structural integrity of euchromatin and heterochromatin, we profiled various chromatin modification marks using CUT&RUN.RNase. We observed that histone protein levels do not change upon RNase A treatment, suggesting that the individual nucleosomes remained stable upon RNA depletion (Supplementary figure 3A). Because histone methylation is a stable covalent modification, we reasoned that if RNA is indeed involved in maintaining chromatin conformations, the disintegration of chromatin domains upon RNA depletion will lead to a decrease in CUT&RUN signals reflecting reduction in the availability of targeted epitopes for antibody binding and/or reduced ability of antibody-tethered pA-MN to access DNA. We performed CUT&RUN.RNase profiling for repressive (H3K27me3 and H3K9me3) and active (H3K4me3) chromatin marks. Surprisingly, genome-wide CUT&RUN signals for the H3K27me3 and H3K9me3 repressive marks decreased dramatically (15 and 8-fold respectively in 5 U RNase A/million cells) upon RNase A treatment (Figure 3A). In contrast, the active H3K4me3 mark showed only a slight decrease (2-fold in 5 U RNase A/million cells) in CUT&RUN signals (Figure 3A). We observed similar effects of RNase A treatment on H3K27me3 and H3K4me3 marks in HeLa cells (Supplementary Figure 3B). These results suggest that the physical presence of RNA is crucial for the structural integrity of both constitutive (H3K9me3) and facultative (H3K27me3) heterochromatic domains.

**Figure 3.**
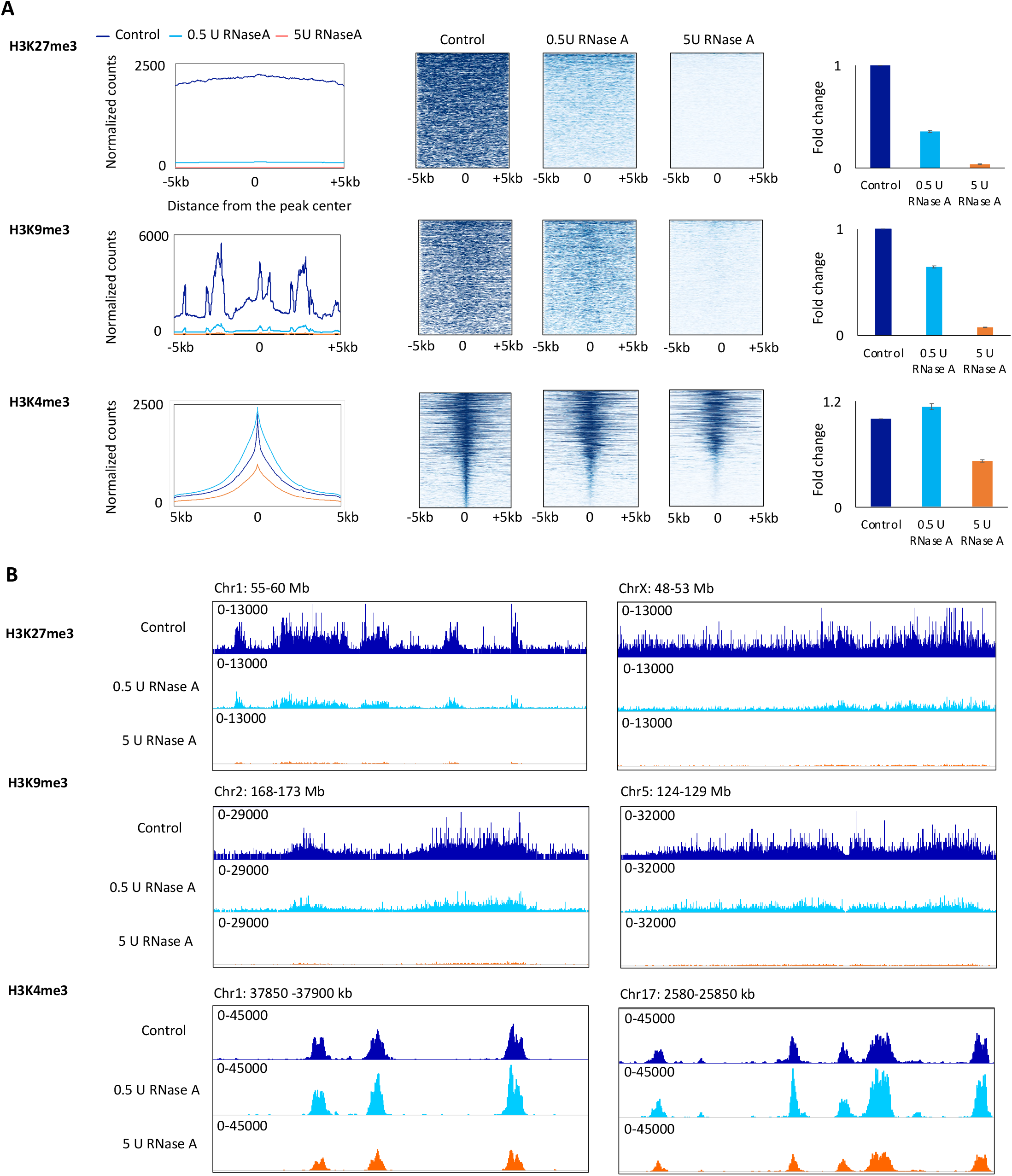
CUT&RUN.RNase detects genome-wide loss of heterochromatin accessibility in RNA-depleted cells. A) Genome-wide H3K27me3, H3K9me3 and H3K4me3 average CUT&RUN signals in control cells and cells treated with two different RNase A concentrations (0.5 and 5 U per million cells) (Right panels). Genome-wide profiles (Left panels) and heatmaps (Center) were generated on a 10 kb region across peak center. B) CUT&RUN.RNase tracks across genomic regions of 5 Mb (H3K27me3 and H3K9me3) or 50 kb (H3K4me3) lengths.

RNA-DNA hybrids including R-loops occupy up to 5% of mammalian genomes and are linked to chromatin compaction at various genomic locations including pericentromeric chromatin (Castellano-Pozo et al., 2013; Sanz et al., 2016; Skourti-Stathaki et al., 2014). Ribonuclease H (RNase H) specifically removes the RNA strand of RNA-DNA hybrids, unlike RNase A, which cleaves single and double-stranded RNA as well the RNA strand of RNA-DNA hybrids. We performed CUT&RUN.RNase by treating cells with RNase H and found that removing RNA-DNA hybrids leads to a slight decrease (~1-3 fold in 5 U RNase H/million cells) in CUT&RUN signals from heterochromatic domains (Supplementary Figure 4). The much higher sensitivity of heterochromatic domains to RNase A than to RNase H implies that R-loops are at most a minor determinant of heterochromatin conformation.

### Cytological decompaction of heterochromatin under CUT&RUN.RNase conditions

To further confirm the disintegration of heterochromatic domains upon RNA depletion, we visualized the cytological appearance of H3K27me3 and H3K9me3 chromatin domains. Patski female murine embryonic kidney fibroblasts are characterized by skewed X-chromosome inactivation and carry an inactive X chromosome (Xi) (Lingenfelter et al., 1998; Yang et al., 2015). The Xi in Patski cells is evident as the brightest H3K27me3 cluster and is associated with the NPM1-stained nucleolus as expected (Yang et al., 2015). The rest of the H3K27me3 puncta were uniformly distributed throughout the nucleus as expected (Figure 4A). The majority of H3K9me3 marked heterochromatin is observed as bright spots at the pericentric regions localized to either the perinucleolar space (co-localized with NPM1 ring) or the nuclear periphery in control cells (Figure 4A). In addition, relatively low levels of H3K9me3 are also present throughout the rest of the nucleus. RNase A treatment led to disappearance or fading of most of the H3K27me3 puncta including that of the Xi (Figure 4A). RNase A treatment also changed perinuclear and perinucleolar H3K9me3 bright distinct spots to uniformly dispersed signals (Figure 4A). Cross-linking the cells prior to RNase A treatment resulted in a complete rescue of both H3K9me3 and H3K27me3 signals (Figure 4B). These results clearly suggest that architectural RNA helps organize H3K27me3 and H3K9me3 marked heterochromatin into highly compact domains at specific locations in the nucleus. In the absence of RNA, these domains become decompacted, lose their native conformation and redistribute into irregular structures.

**Figure 4.**
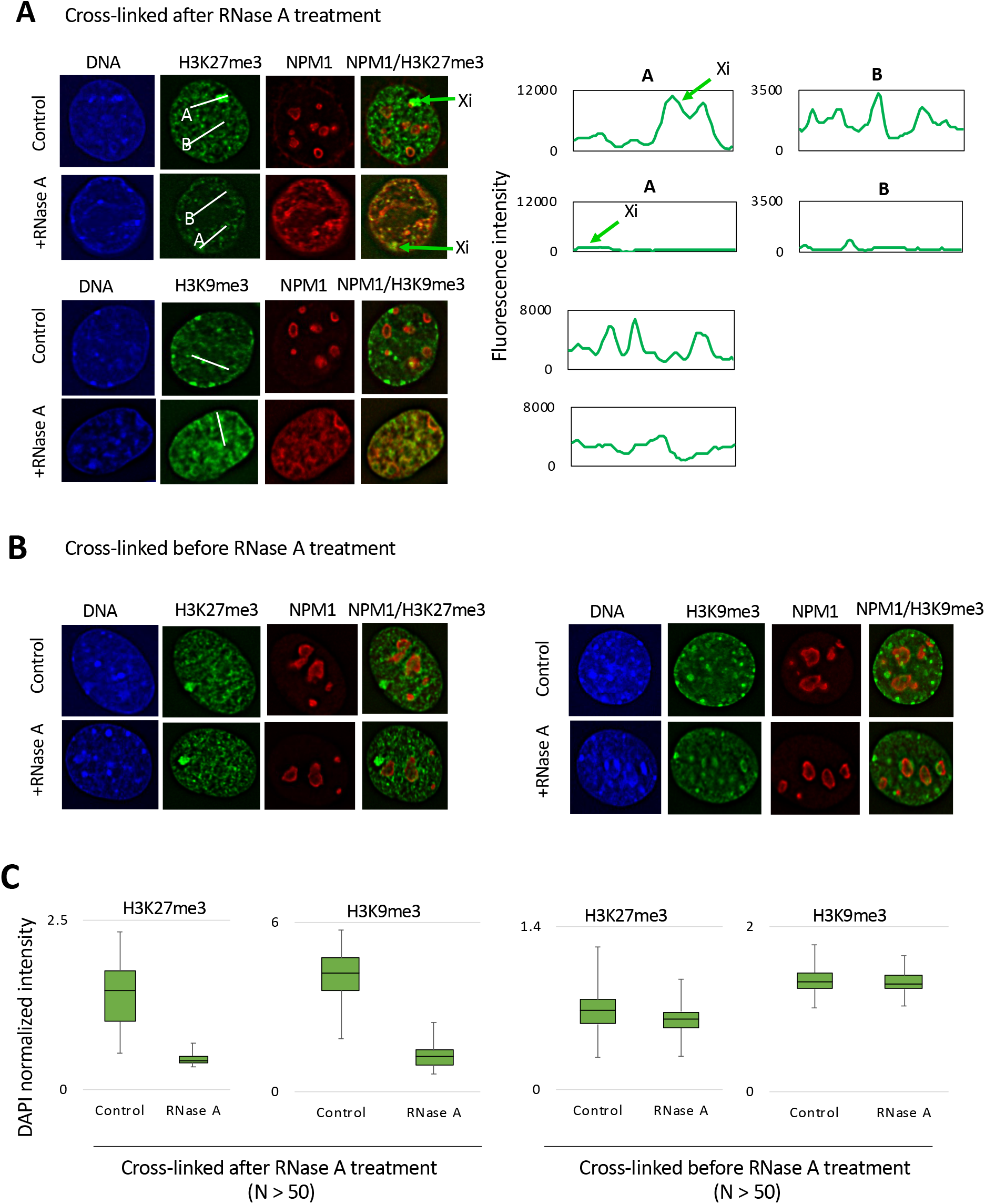
Decompaction of heterochromatin under CUT&RUN.RNase conditions. A) Patski cells treated with RNase A, cross-linked and stained with NPM1, DAPI and H3K27me3 or H3K9me3. The intensity profiles shown in the right panels were plotted across the white lines drawn on the images shown in the left panels. B) Patski cells cross-linked, RNase A treated and stained with NPM1, DAPI and H3K27me3 or H3K9me3 C) The intensity of H3K27me3 or H3K9me3 signals normalized to the DAPI intensity for the indicated experimental conditions.

### *Firre* lncRNA deletion disrupts a subset of H3K27me3 domains

CUT&RUN.RNase suggests that global depletion of RNA results in genome-wide loss of heterochromatin integrity. To establish the specificity and biological relevance of heterochromatin stabilization by RNA, we investigated the effect on heterochromatin organization of deleting a single lncRNA. *Firre*, a widely distributed lncRNA in the nucleus, is transcribed from a macrosatellite repeat locus on X chromosome and establishes contacts with multiple autosomes (Hacisuleyman et al., 2016; Yang et al., 2015). *Firre* depletion leads to a decrease in H3K27me3 levels on the Xi as well as change in the expression of several autosomal genes (Bergmann et al., 2015; Rinn et al., 2007; Yang et al., 2015). Therefore, we wondered whether the genome-wide loss of heterochromatin integrity that we observed using RNase A would be locally recapitulated on the Xi and specific autosomal sites with loss of the *Firre* lncRNA. We utilized H3K27me3 chromatin on the Xi as a positive control for detecting the effect of *Firre* depletion on heterochromatin domains. The *Firre* locus has been deleted using allele-specific CRISPR/Cas9 editing in *ΔFirre^Xa^*, a derivative of Patski-WT (He et al., 2019). Deletion of the *Firre* locus from the active X-chromosome (Xa) results in undetectable levels of *Firre* transcripts and loss of H3K27me3 from the inactive X-chromosome (He et al., 2019). We performed H3K27me3 and H3K4me3 CUT&RUN in both WT and *ΔFirre^Xa^* cells. A comparison of CUT&RUN signals between the WT and *ΔFirre^Xa^* revealed a 20 percent decrease in H3K27me3 signals in *Firre* deleted cells. The reduction in H3K27me3 signals in *ΔFirre^Xa^* was ~10 times lower than what was observed upon RNase A treatment (5 U/million cells) suggesting that *Firre* contributes to a small subset of all RNA-H3K27me3 chromatin interactions. Global H3K4me3 CUT&RUN signals showed no significant differences between the WT and *ΔFirre^Xa^* (Figure 5A).

**Figure 5.**
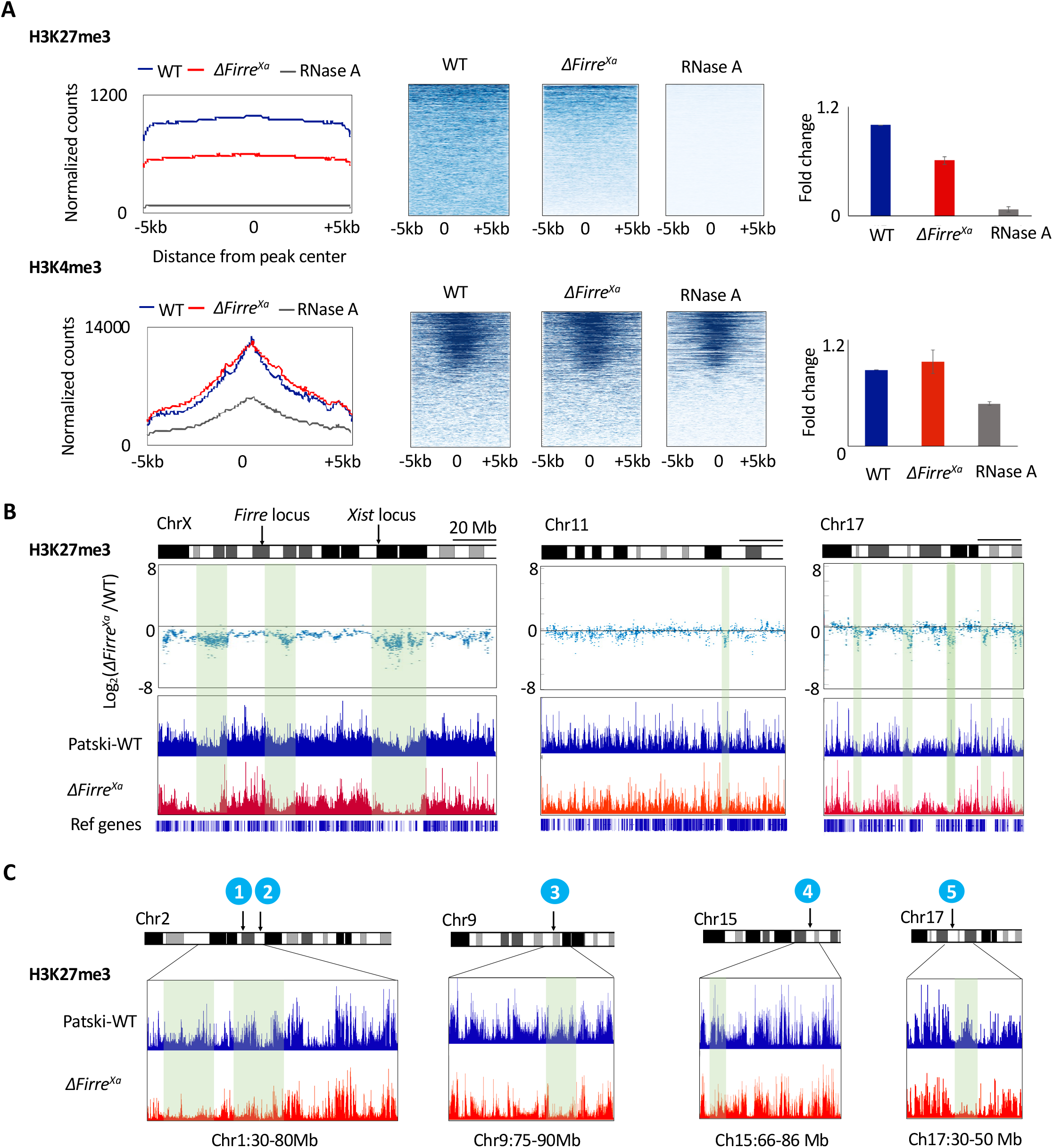
*Firre* depletion reduces H3K27me3 CUT&RUN signal at a subset of domains. A) Genome-wide H3K27me3 and H3K4me3 average CUT&RUN signals in WT, *ΔFirre^Xa^* and RNase A treated WT (5 U per million cells). Genome-wide profiles (Left) and heatmaps (Middle panels) were generated on a 10 kb region across peak center. Fold changes are shown relative to WT (Right) B) Log2 ratio of WT and *ΔFirre^Xa^* H3K27me3 signals on ChrX, Chr11 and Chr17 H3K27me3 domains (Top panels). Autoscaled H3K27me3 signal tracks on ChrX, Chr11 and Chr17 (Bottom panels). NCBI RefSeq genes are displayed below H3K27me3 signal tracks for each ChrX, Chr11 and Chr17. C) Autoscaled H3K27me3 signal tracks on chromosomal regions of indicated lengths spanning *Firre* interacting sites.

To identify the regions affected by *Firre* deletion, we plotted Log2 ratios of WT and *ΔFirre^Xa^* H3K27me3 CUT&RUN signals on individual chromosomes. On Chromosome X, we identified three large regions spanning tens of megabases that showed loss of H3K27me3 signal in *ΔFirre^Xa^* (Figure 5B). Two out of three regions flank the *Firre* locus while the third region is located adjacent to the *Xist* locus. Interestingly, we also noticed loss of CUT&RUN signals at megabase-sized H3K27me3 domains on several autosomes (Figure 5B and 5C). We asked if affected regions include five *Firre* interacting autosomal loci that were previously identified using RNA antisense purification (Hacisuleymanet al., 2014). We found that indeed each of these five *Firre* interacting loci were located near regions that showed the loss of H3K27me3 CUT&RUN signals in *ΔFirre^Xa^*. Our results with *ΔFirre^Xa^* confirm the biological significance of our CUT&RUN.RNase approach and suggest that the global effect of RNA depletion that we observed in CUT&RUN.RNase result primarily from disruption of specific interactions between various lncRNAs and heterochromatin. Moreover, the similar effects of *Firre* deletion in vivo and RNase A treatment on heterochromatin domains in situ further suggests that the mechanism of action of lncRNAs on X-chromosome inactivation is applicable to the entire genome.

## Discussion

Mature RNA is abundant in the nucleus and is often associated with chromatin (Bell et al., 2018; Pandey et al., 2008; Rodríguez-Campos and Azorín, 2007; West et al., 2014). However, understanding the contribution of RNA to chromatin organization resulting from direct RNA-chromatin contacts has remained a challenge using current genomic technologies. The CUT&RUN chromatin profiling method is performed in situ, which allows for probing the chromatin environment around a given epitope at high resolution (Skene and Henikoff, 2017). The use of RNase A digestion under gentle in situ conditions has allowed us to dissect the contribution of RNA due to its physical presence on chromatin from its indirect regulatory role in chromatin, for example via recruitment of chromatin modifiers. Combining CUT&RUN with RNase digestion provides a unique opportunity to decipher the contribution of the presence of RNA in chromatin organization at high resolution. As a proof of principle, using CUT&RUN.RNase we demonstrated that interactions of nucleolar protein NPM1 with pericentric heterochromatin at the nucleolar periphery are dependent on the presence of RNA.

The architectural protein CTCF insulates adjacent active and repressed chromatin domains and also mediates 3-D contacts at several developmentally regulated genomic loci (Cuddapah et al., 2009; Fu et al., 2008; Narendra et al., 2015; Phillips and Corces, 2009) Owens et al., 2019, Clarkson et al., 2019). In addition to precisely mapping direct transcription factor footprints, CUT&RUN also probes the local and 3-D chromatin environments around CTCF binding sites (Skene and Henikoff, 2017). CTCF CUT&RUN.RNase revealed that the intact RNA is required for maintaining the chromatin environment around CTCF likely by facilitating local chromatin compaction, such that loss of RNA reduces CTCF-tethered MNase cleavage of the linker DNA around the adjacent nucleosomes.

The majority of heterochromatin is spread across large regions that are condensed to form spatially compact domains. In case of X inactivation, H3K27me3 heterochromatin is spread across the length of entire X-chromosome, which then forms a highly compact structure called the Barr body near the nucleolus (Engreitz et al., 2013; Lee, 2003; Pinheiro and Heard, 2017). Pericentric H3K9me3 heterochromatin is formed across a several megabase long highly repetitive satellite DNA array on every chromosome. In mouse, pericentric heterochromatin is mostly organized around nucleoli (Guenatri et al., 2004). Based on in vitro studies, a specific hierarchical higher order structure was believed to be the basis of heterochromatin compaction (Tremethick, 2007). However visualization of chromatin at nucleosome resolution has revealed chromatin to be a disordered chain of nucleosomes where euchromatin and heterochromatin differ from each other in terms of their local density (Ou et al., 2017). Our results reveal that the presence of RNA is essential for the integrity and compactness of both constitutive and facultative heterochromatic domains. The architectural role of RNA that CUT&RUN.RNase revealed may be a general property of chromatin-associated RNA, may represent multiple specific RNA-chromatin interactions or a combination of both.

There have been several reports of specific cis and trans RNA-chromatin interactions (Engreitz et al., 2013; Pandey et al., 2008; Pinheiro and Heard, 2017; Rego et al., 2008; Rinn et al., 2007). *Firre* lncRNA, apart from its role in maintaining H3K27me3 chromatin state on the inactive X, also makes contacts with several autosomes. Interestingly, mouse *Firre* contains 16 copies of a repeating domain (Hacisuleyman et al., 2014), which might facilitate establishment of anchor points within discrete 3-D proximal spaces. Thus, *Firre* is a suitable candidate lncRNA to investigate the possibility that a specific lncRNA might be involved in heterochromatin organization at multiple chromosomes. Our results show that in addition to maintaining heterochromatin on the X-chromosome as previously reported, deletion of *Firre* leads to disruption of H3K27me3 domains on multiple autosomes in vivo. Loss of signals on H3K27me3 domains in *Firre*-deleted cells spanned several megabase regions suggesting that, similar to their role in maintaining Xi heterochromatin, lncRNAs might play a much broader role in heterochromatin organization.

## Material and methods

### Cell lines and antibodies

Mouse embryonic kidney fibroblast cell lines Patski wild-type and ΔFirreXa (Fang H et al., 2019, (Lingenfelter et al., 1998), were grown in DMEM media containing 13% fetal bovine serum. Antibodies used for CUT&RUN.RNase, Western and immunofluorescence were H3K27me3 (Cat# 9733, Cell Signalling Technology), H3K9me3 (Cat# ab8898, Abcam), H3K4me3 (Cat# 39060, Active Motif), CTCF (Cat # 07-729, Millipore Sigma), NPM1, (Cat # ab10530, Abcam), Tubulin (Cat # T5168, Sigma Aldrich) and Histone H3 (Cat # 1791, Abcam).

### CUT&RUN.RNase

Cells were bound to Concanavalin A beads as described previously, beads were resuspended in HCMD buffer (20 mM HEPES pH 7.5, 0.1 mM CaCl2, 3mM MgCl2, 100 mM KCl, 0.05% digitonin, Protease inhibitor) and divided into two equal aliquots. To one aliquot, RNase A or RNase H was added at a concentration of 5 U per million cells (unless stated otherwise). The cells with or without RNase A were incubated for 90 min at room temperature followed by 3 washes with wash buffer (20 mM HEPES, pH 7.5, 150 mM NaCl, 0.5 mM spermidine, 0.05% digitonin, Protease inhibitor). Equal amounts of antibodies were added to both control and RNase A treated cells. The remainder of the CUT&RUN protocol was performed as described previously (Skene and Henikoff, 2017).

For cross-linking CUT&RUN.RNAse, cells were cross-linked with 1% formaldehyde for 10 min at room temperature followed by CUT&RUN.RNase as described above. After adding the stop buffer (170 mM NaCl, 20 mM EGTA, 0.05% Digitonin, 20 µg/ml glycogen, 25 µg/ml RNase A, 5 pg/ml

S. cerevisiae fragmented nucleosomal DNA), the samples were incubated on a nutator for 2 hrs at 40C. The supernatant was separated from the beads and de-crosslinked overnight at 650C. De-crosslinked samples were then subjected to DNA extraction as described previously.

### Sequencing and Data processing

Library preparation from CUT&RUN DNA and subsequent sequencing on the Illumina HiSeq 2500 platform were carried out as described previously (Skene and Henikoff, 2017) to obtain 25×25 paired end reads. Paired-end fragments were mapped to the sacCer3/V64 and mouse mm10 genomes using Bowtie2 version 2.2.5 with options: --local --very-sensitive-local --no-unal --no-mixed --no-discordant to each mouse CUT&RUN reaction. We mapped the pairedend reads to both mouse and yeast genomes and calibrated datasets by dividing the number of mapped mouse fragments by the number of fragments mapped to the yeast genome. Average profiles of NPM1 on major satellites were generated by mapping NPM1 CUT&RUN-generated fragments to a concatenated hexamer of a 234 bp major satellite repeat unit. The CTCF profiles were generated across a 5 kb region surrounding annotated CTCF sites (Encode Project). CTCF V-plots were generated as described previously (Henikoff et al., 2011). Human direct and indirect CTCF sites used for mapping HeLa CTCF CUT&RUN.RNase data have been described previously (Skene and Henikoff, 2017).

### Immunofluorescence

Cells were grown overnight in chambered slides (Cat# PEZGS0416, Millipore) at a final density of ~300,000 cells per well. A 400 µl volume of HCMD buffer (20 mM HEPES pH 7.5, 0.1 mM CaCl2, 3mM MgCl2, 100 mM KCl, 0.05% digitonin, Protease inhibitor) was added per well. A series of RNase A concentrations was tested on Patski cells to determine the concentration at which clear effects of RNase A digestion on chromatin marks were observed without significantly damaging the overall nuclear structure (as determined by DAPI staining). For the experiments presented here, 12.5 U of RNase A was added per well (~300,000 Patski cells). Both control and RNase A containing slides were incubated for 90 min and immediately fixed with 4% formaldehyde. Cells were incubated in phosphate-buffered saline + 0.05% Triton-X + 1% bovine serum albumin solution for 30 min. Cells were then incubated in primary antibodies overnight, washed 5 times with PBST, incubated in secondary antibodies for 45 min, washed 5 times with PBST and mounted using a glycerol-containing DAPI solution. Images were captured using a DeltaVision Elite microscope, and quantification was performed using Fiji (ImageJ) software.

## Funding

This work was supported by NIH grants HG010492 (SH) and GM131745 (CMD) and by the NIH Common Fund 4D Nucleome Program DK107979.

## Acknowledgments

We thank Christine Codomo for preparing Illumina sequencing libraries, Jorja Henikoff for bioinformatics, the Fred Hutch Genomics Shared Resource for sequencing and the Fred Hutch Cellular Imaging Shared Resource for imaging. We also thank Henikoff lab members for helpful discussions.

---

**Supplementary Figure 1.**
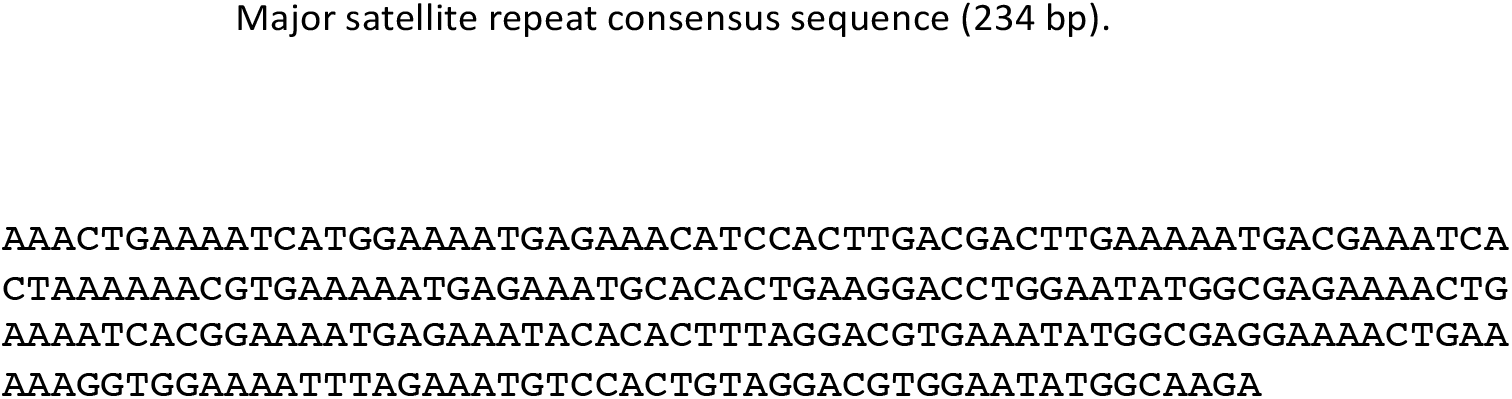
The consensus major satellite sequence

**Supplementary Figure 2.**
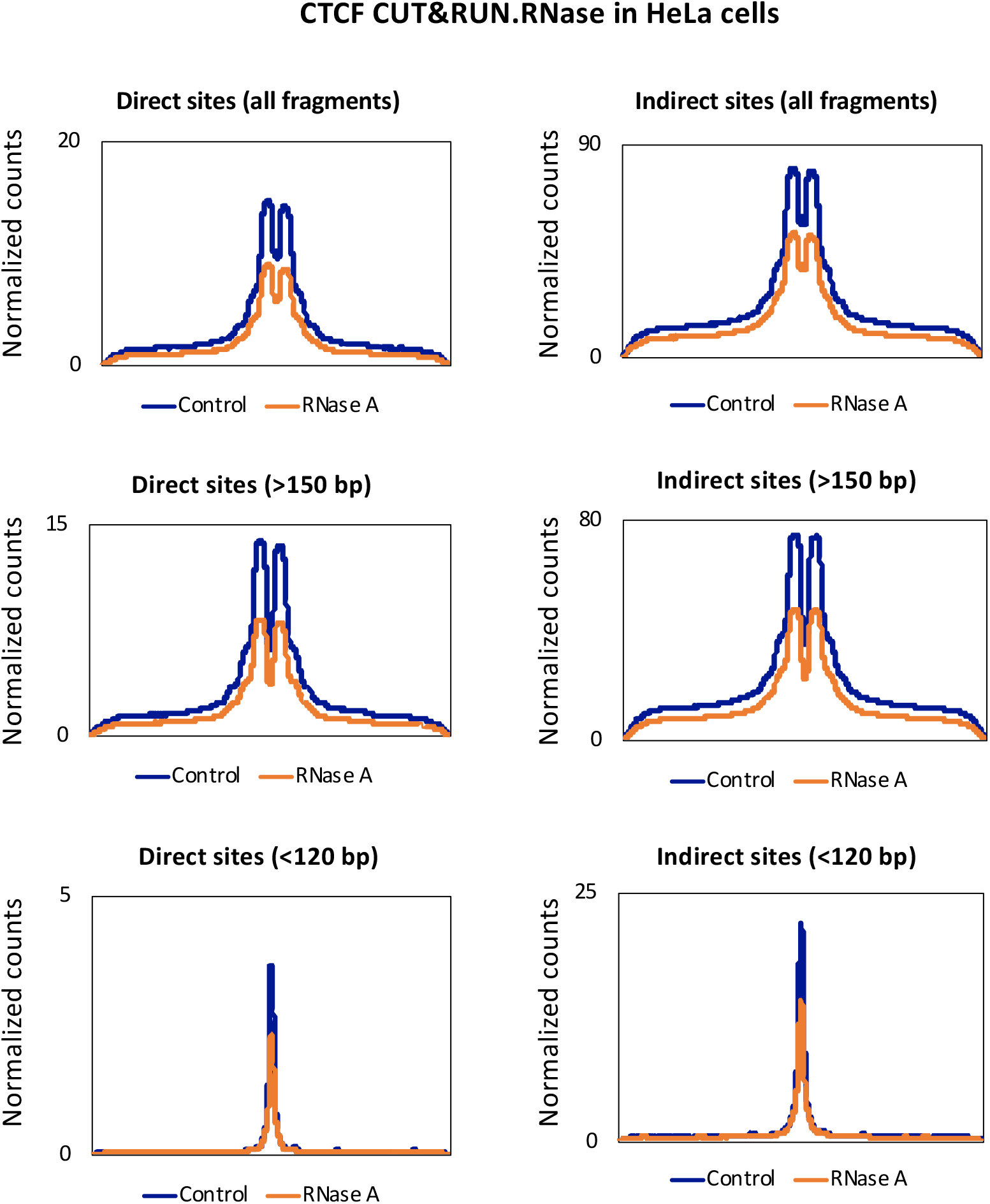
CTCF CUT&RUN.RNAse signals for all, small (< 120 bp) and large (> 150 bp) fragments in control and RNase A treated HeLa cells (5 U RNase A/million cells). Genome-wide average profiles (Left) were generated on a 5 kb region across annotated CTCF sites.

**Supplementary Figure 3.**
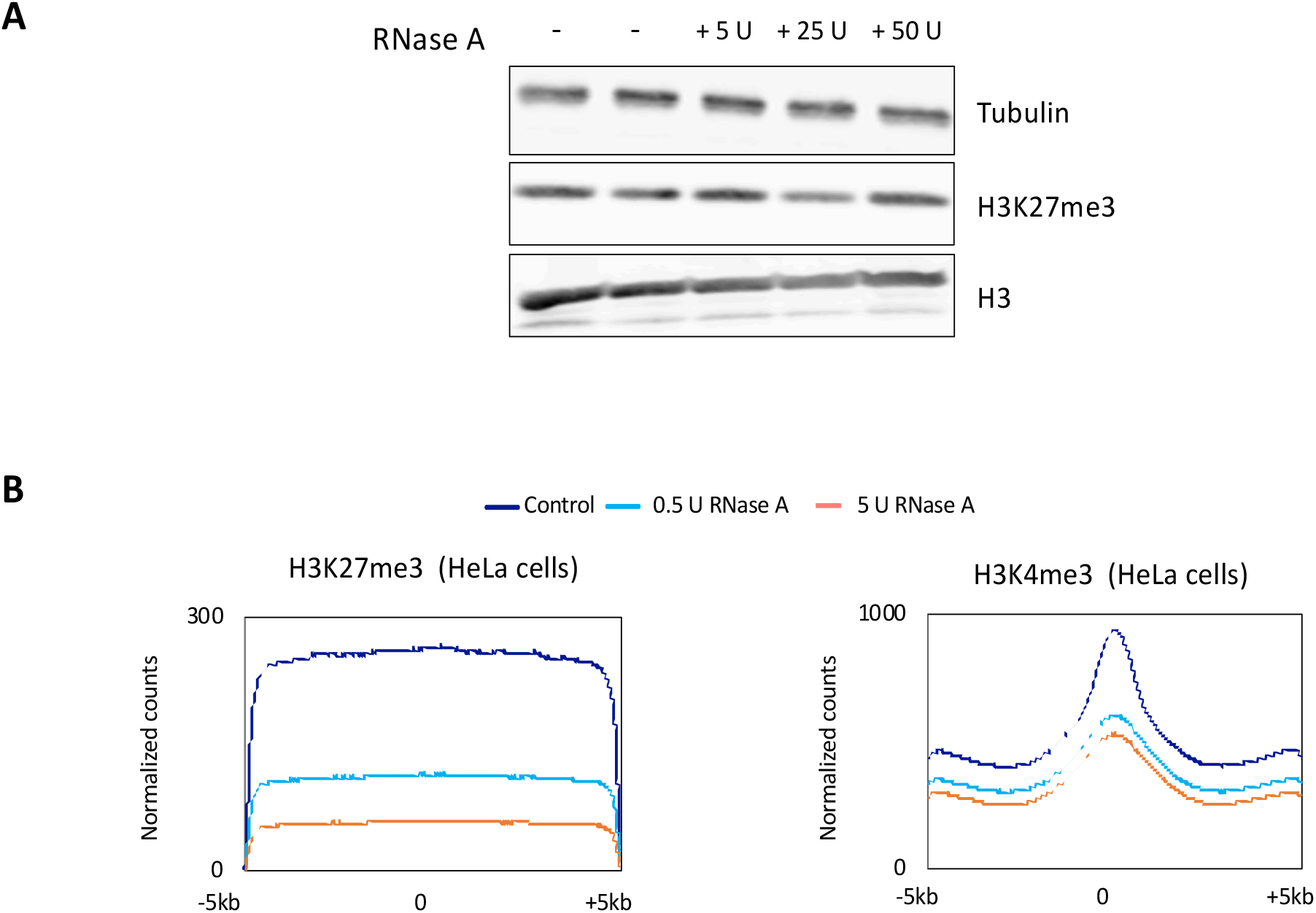
A) Western blot assays showing Tubulin, H3K27me3 and H3 protein levels. B) H3K27me3 and H3K4me3 CUT&RUN.RNase in HeLa cells. Genome-wide average profiles were generated on a 10 kb region across the peak center.

**Supplementary Figure S4.**
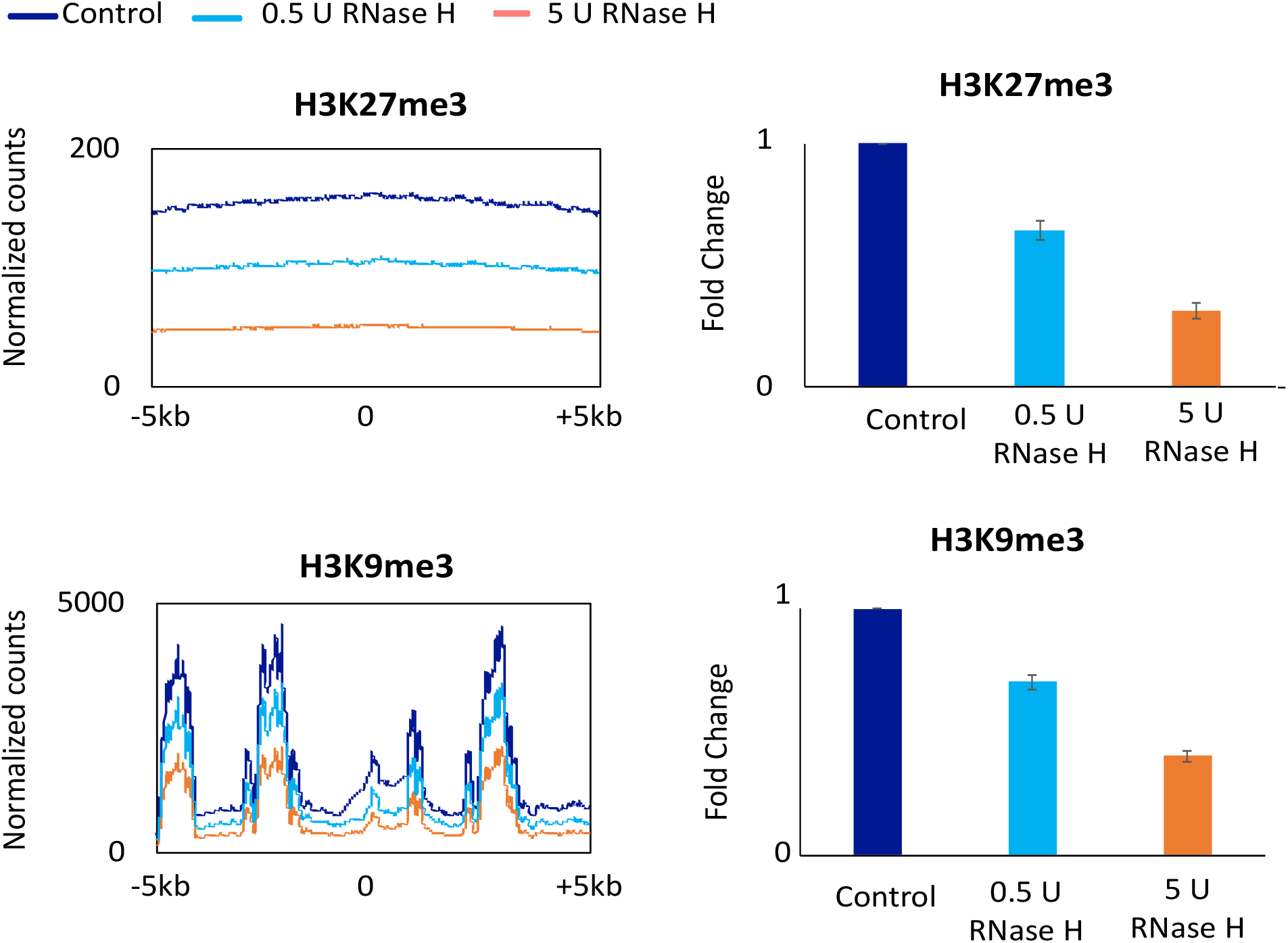
H3K27me3 and H3K9me3 CUT&RUN.RNase in control cells and cells treated with two different RNase H concentrations (0.5 and 5 U per million Patski cells). Genome-wide average profiles (Left) were generated on a 10 kb region across the peak center. Genome-wide H3K27me3, H3K9me3 and H3K4me3 average CUT&RUN signals in control and RNase H treated cells (Right).

## References

Almouzni, G., and Probst, A.V. (2011). Heterochromatin maintenance and establishment: lessons from the mouse pericentromere. Nucleus 2, 332–338.

Anders S. HansenTsung-Han S. Hsieh, Claudia Cattoglio, Iryna Pustova, Xavier Darzacq, Robert Tjian (2019). An RNA-binding region regulates CTCF clustering and chromatin looping. biorxiv

Bannister, A.J., Zegerman, P., Partridge, J.F., Miska, E.A., Thomas, J.O., Allshire, R.C., and Kouzarides, T. (2001). Selective recognition of methylated lysine 9 on histone H3 by the HP1 chromo domain. Nature 410, 120–124.

Bártová, E., Krejcí, J., Harnicarová, A., Galiová, G., and Kozubek, S. (2008). Histone modifications and nuclear architecture: a review. J Histochem Cytochem 56, 711–721.

Barutcu, A.R., Blencowe, B.J., and Rinn, J.L. (2019). Differential contribution of steady-state RNA and active transcription in chromatin organization. EMBO Rep, e48068.

Bell, J.C., Jukam, D., Teran, N.A., Risca, V.I., Smith, O.K., Johnson, W.L., Skotheim, J.M., Greenleaf, W.J., and Straight, A.F. (2018). Chromatin-associated RNA sequencing (ChAR-seq) maps genome-wide RNA-to-DNA contacts. Elife 7.

Bergmann, J.H., Li, J., Eckersley-Maslin, M.A., Rigo, F., Freier, S.M., and Spector, D.L. (2015). Regulation of the ESC transcriptome by nuclear long noncoding RNAs. Genome Res 25, 1336–1346.

Boisvert, F.M., van Koningsbruggen, S., Navascués, J., and Lamond, A.I. (2007). The multifunctional nucleolus. Nat Rev Mol Cell Biol 8, 574–585.

Burke, B., and Stewart, C.L. (2014). Functional architecture of the cell’s nucleus in development, aging, and disease. Curr Top Dev Biol 109, 1–52.

Castellano-Pozo, M., Santos-Pereira, J.M., Rondón, A.G., Barroso, S., Andújar, E., Pérez-Alegre, M., García-Muse, T., and Aguilera, A. (2013). R loops are linked to histone H3 S10 phosphorylation and chromatin condensation. Mol Cell 52, 583–590.

Cheutin, T., and Cavalli, G. (2012). Progressive polycomb assembly on H3K27me3 compartments generates polycomb bodies with developmentally regulated motion. PLoS Genet 8, e1002465.

Christopher T. Clarkson, Emma A. Deeks, Ralph Samarista, Hulkar Mamayusupova, Victor B. Zhurkin, Vladimir B. Teif (2019) CTCF-dependent chromatin boundaries formed by asymmetric nucleosome arrays with decreased linker length. biorxiv

Cuddapah, S., Jothi, R., Schones, D.E., Roh, T.Y., Cui, K., and Zhao, K. (2009). Global analysis of the insulator binding protein CTCF in chromatin barrier regions reveals demarcation of active and repressive domains. Genome Res 19, 24–32.

Engreitz, J.M., Ollikainen, N., and Guttman, M. (2016). Long non-coding RNAs: spatial amplifiers that control nuclear structure and gene expression. Nat Rev Mol Cell Biol 17, 756–770.

Engreitz, J.M., Pandya-Jones, A., McDonel, P., Shishkin, A., Sirokman, K., Surka, C., Kadri, S., Xing, J., Goren, A., Lander, E.S., et al. (2013). The Xist lncRNA exploits three-dimensional genome architecture to spread across the X chromosome. Science 341, 1237973.

Fu, Y., Sinha, M., Peterson, C.L., and Weng, Z. (2008). The insulator binding protein CTCF positions 20 nucleosomes around its binding sites across the human genome. PLoS Genet 4, e1000138.

Grewal, S.I. (2010). RNAi-dependent formation of heterochromatin and its diverse functions. Curr Opin Genet Dev 20, 134–141.

Guenatri, M., Bailly, D., Maison, C., and Almouzni, G. (2004). Mouse centric and pericentric satellite repeats form distinct functional heterochromatin. J Cell Biol 166, 493–505.

Hacisuleyman, E., Goff, L.A., Trapnell, C., Williams, A., Henao-Mejia, J., Sun, L., McClanahan, P., Hendrickson, D.G., Sauvageau, M., Kelley, D.R., et al. (2014). Topological organization of multichromosomal regions by the long intergenic noncoding RNA Firre. Nat Struct Mol Biol 21, 198–206.

Hacisuleyman, E., Shukla, C.J., Weiner, C.L., and Rinn, J.L. (2016). Function and evolution of local repeats in the Firre locus. Nat Commun 7, 11021.

Heitz E. Das Heterochromatin der Moose. Jahrb Wiss Botanik. 1928;69:762–818

He Fang, Giancarlo Bonora, Jordan P. Lewandowski, Jitendra Thakur, Galina N. Filippova, Steven Henikoff, Jay Shendure, Zhijun Duan, John L. Rinn, Xinxian Deng, William S. Noble, Christine M. Disteche (2019). Trans- and cis-acting effects of the lncRNA Firre on epigenetic and structural features of the inactive X chromosome. biorxiv

Henikoff, J.G., Belsky, J.A., Krassovsky, K., MacAlpine, D.M., and Henikoff, S. (2011). Epigenome characterization at single base-pair resolution. Proc Natl Acad Sci U S A 108, 18318–18323.

Holmberg Olausson, K., Nistér, M., and Lindström, M.S. (2014). Loss of nucleolar histone chaperone NPM1 triggers rearrangement of heterochromatin and synergizes with a deficiency in DNA methyltransferase DNMT3A to drive ribosomal DNA transcription. J Biol Chem 289, 34601–34619.

Lachner, M., O’Carroll, D., Rea, S., Mechtler, K., and Jenuwein, T. (2001). Methylation of histone H3 lysine 9 creates a binding site for HP1 proteins. Nature 410, 116–120.

Lee, J.T. (2003). X-chromosome inactivation: a multi-disciplinary approach. Semin Cell Dev Biol 14, 311–312.

Lindström, M.S. (2011). NPM1/B23: A Multifunctional Chaperone in Ribosome Biogenesis and Chromatin Remodeling. Biochem Res Int 2011, 195209.

Lingenfelter, P.A., Adler, D.A., Poslinski, D., Thomas, S., Elliott, R.W., Chapman, V.M., and Disteche, C.M. (1998). Escape from X inactivation of Smcx is preceded by silencing during mouse development. Nat Genet 18, 212–213.

Maison, C., Bailly, D., Peters, A.H., Quivy, J.P., Roche, D., Taddei, A., Lachner, M., Jenuwein, T., and Almouzni, G. (2002). Higher-order structure in pericentric heterochromatin involves a distinct pattern of histone modification and an RNA component. Nat Genet 30, 329–334.

Mitrea, D.M., Cika, J.A., Guy, C.S., Ban, D., Banerjee, P.R., Stanley, C.B., Nourse, A., Deniz, A.A., and Kriwacki, R.W. (2016). Nucleophosmin integrates within the nucleolus via multi-modal interactions with proteins displaying R-rich linear motifs and rRNA. Elife 5.

Murano, K., Okuwaki, M., Hisaoka, M., and Nagata, K. (2008). Transcription regulation of the rRNA gene by a multifunctional nucleolar protein, B23/nucleophosmin, through its histone chaperone activity. Mol Cell Biol 28, 3114–3126.

Nagano, T., Mitchell, J.A., Sanz, L.A., Pauler, F.M., Ferguson-Smith, A.C., Feil, R., and Fraser, P. (2008). The Air non-coding RNA epigenetically silences transcription by targeting G9a to chromatin. Science 322, 1717–1720.

Narendra, V., Rocha, P.P., An, D., Raviram, R., Skok, J.A., Mazzoni, E.O., and Reinberg, D. (2015). CTCF establishes discrete functional chromatin domains at the Hox clusters during differentiation. Science 347, 1017–1021.

Nick Owens, Thaleia Papadopoulou, Nicola Festuccia, Alexandra Tachtsidi, Inma Gonzalez, Agnes Dubois, Sandrine Vandormael-Pournin, Elphege P Nora, Benoit G Bruneau, Michel Cohen-Tannoudji, Pablo Navarro (2019) CTCF confers local nucleosome resiliency after DNA replication and during mitosis. biorxiv

Nickerson, J.A., Krochmalnic, G., Wan, K.M., and Penman, S. (1989). Chromatin architecture and nuclear RNA. Proc Natl Acad Sci U S A 86, 177–181.

Ou, H.D., Phan, S., Deerinck, T.J., Thor, A., Ellisman, M.H., and O’Shea, C.C. (2017). ChromEMT: Visualizing 3D chromatin structure and compaction in interphase and mitotic cells. Science 357.

Pandey, R.R., Mondal, T., Mohammad, F., Enroth, S., Redrup, L., Komorowski, J., Nagano, T., Mancini-Dinardo, D., and Kanduri, C. (2008). Kcnq1ot1 antisense noncoding RNA mediates lineage-specific transcriptional silencing through chromatin-level regulation. Mol Cell 32, 232–246.

Park, P.J. (2009). ChIP-seq: advantages and challenges of a maturing technology. Nat Rev Genet 10, 669–680.

Phillips, J.E., and Corces, V.G. (2009). CTCF: master weaver of the genome. Cell 137, 1194–1211.

Pinheiro, I., and Heard, E. (2017). X chromosome inactivation: new players in the initiation of gene silencing. F1000Res 6.

Politz, J.C., Scalzo, D., and Groudine, M. (2013). Something silent this way forms: the functional organization of the repressive nuclear compartment. Annu Rev Cell Dev Biol 29, 241–270.

Prensner, J.R., Iyer, M.K., Sahu, A., Asangani, I.A., Cao, Q., Patel, L., Vergara, I.A., Davicioni, E., Erho, N., Ghadessi, M., et al. (2013). The long noncoding RNA SChLAP1 promotes aggressive prostate cancer and antagonizes the SWI/SNF complex. Nat Genet 45, 1392–1398.

Rasband, W.S., ImageJ, U. S. National Institutes of Health, Bethesda, Maryland, USA, https://imagej.nih.gov/ij/, 1997–2018.

Ricardo Saldana-Meyer, Javier Rodriguez-Hernaez, Mayilaadumveettil Nishana, Karina Jacome-Lopez, Elphege P. Nora, Benoit G. Bruneau, Mayra Furlan-Magaril, Jane Skok, Danny Reinberg (2019) RNA interactions with CTCF are essential for its proper function. biorxiv

Rego, A., Sinclair, P.B., Tao, W., Kireev, I., and Belmont, A.S. (2008). The facultative heterochromatin of the inactive X chromosome has a distinctive condensed ultrastructure. J Cell Sci 121, 1119–1127.

Rinn, J.L., Kertesz, M., Wang, J.K., Squazzo, S.L., Xu, X., Brug-mann, S.A., Goodnough, L.H., Helms, J.A., Farnham, P.J., Segal, E., et al. (2007). Functional demarcation of active and silent chromatin domains in human HOX loci by noncoding RNAs. Cell 129, 1311–1323.

Rodríguez-Campos, A., and Azorín, F. (2007). RNA is an integral component of chromatin that contributes to its structural organization. PLoS One 2, e1182.

Sanz, L.A., Hartono, S.R., Lim, Y.W., Steyaert, S., Rajpurkar, A., Ginno, P.A., Xu, X., and Chédin, F. (2016). Prevalent, Dynamic, and Conserved R-Loop Structures Associate with Specific Epigenomic Signatures in Mammals. Mol Cell 63, 167–178.

Skene, P.J., and Henikoff, S. (2017). An efficient targeted nuclease strategy for high-resolution mapping of DNA binding sites. Elife 6.

Skourti-Stathaki, K., Kamieniarz-Gdula, K., and Proudfoot, N.J. (2014). R-loops induce repressive chromatin marks over mammalian gene terminators. Nature 516, 436–439.

Tremethick, D.J. (2007). Higher-order structures of chromatin: the elusive 30 nm fiber. Cell 128, 651–654.

Verdel, A., Jia, S., Gerber, S., Sugiyama, T., Gygi, S., Grewal, S.I., and Moazed, D. (2004). RNAi-mediated targeting of heterochromatin by the RITS complex. Science 303, 672–676.

West, J.A., Davis, C.P., Sunwoo, H., Simon, M.D., Sadreyev, R.I., Wang, P.I., Tolstorukov, M.Y., and Kingston, R.E. (2014). The long noncoding RNAs NEAT1 and MALAT1 bind active chromatin sites. Mol Cell 55, 791–802.

Yang, F., Deng, X., Ma, W., Berletch, J.B., Rabaia, N., Wei, G., Moore, J.M., Filippova, G.N., Xu, J., Liu, Y., et al. (2015). The lncRNA Firre anchors the inactive X chromosome to the nucleolus by binding CTCF and maintains H3K27me3 methylation. Genome Biol 16, 52.

Yap, K.L., Li, S., Muñoz-Cabello, A.M., Raguz, S., Zeng, L., Mujtaba, S., Gil, J., Walsh, M.J., and Zhou, M.M. (2010). Molecular interplay of the noncoding RNA ANRIL and methylated histone H3 lysine 27 by polycomb CBX7 in transcriptional silencing of INK4a. Mol Cell 38, 662–674.

Zhao, J., Sun, B.K., Erwin, J.A., Song, J.J., and Lee, J.T. (2008). Polycomb proteins targeted by a short repeat RNA to the mouse X chromosome. Science 322, 750–756.

Zhou, K., Gaullier, G., and Luger, K. (2019). Nucleosome structure and dynamics are coming of age. Nat Struct Mol Biol 26, 3–13.

